# Vapor cannabis exposure generationally affects male reproductive functions in mice

**DOI:** 10.1101/2021.08.22.457271

**Authors:** Mingxin Shi, Esther M. Langholt, Logan C. Butler, Madeleine E. Harvey, Emma C. Wheeler, Liang Zhao, James A. MacLean, Yeongseok Oh, Emily Sabrowsky, Sue Yu, Shane Watson, Jon F. Davis, Kanako Hayashi

**Author notes:** Correspondence: Kanako Hayashi, School of Molecular Biosciences, Center for Reproductive Biology, Washington State University, 1770 NE Stadium Way, Pullman, WA, 99164, USA. MS and EML contributed equally to this work.

## Abstract

This study was performed to examine whether vapor exposure to cannabis plant matter negatively impacts male reproductive functions and testis development in mice. Adult CD-1 male mice (F0) were exposed to air (control) or 200 mg of vaporized cannabis plant matter 3x/day over a 10 day period. Subsequently, F0 males were bred with drug naïve CD-1 females to generate F1 males, and F1 offspring were used to generate F2 males. Cannabis vapor exposure decreased sperm count and/or motility in F0 and F1 males and disrupted the progression of germ cell development, as morphometric analyses exhibited an abnormal distribution of the stages of spermatogenesis in F0 males. Although plasma levels of testosterone were not affected by cannabis exposure in any ages or generations of males, dysregulated steroidogenic enzymes, *Cyp11a1* and *Cyp19a1*, were observed in F0 testis. In the neonatal testis from F1 males, while apoptosis was not altered, DNA damage and DNMT1, but not DNMT3A and DNMT3B, were increased in germ cells following cannabis exposure. In contrast, the alterations of DNA damage and DNMT1 expression were not observed in F2 neonatal males. These results suggest that cannabis vapor exposure generationally affects male reproductive functions, probably due to disruption of spermatogenesis in the developing testis.

**GRAPHICAL ABSTRACT:** 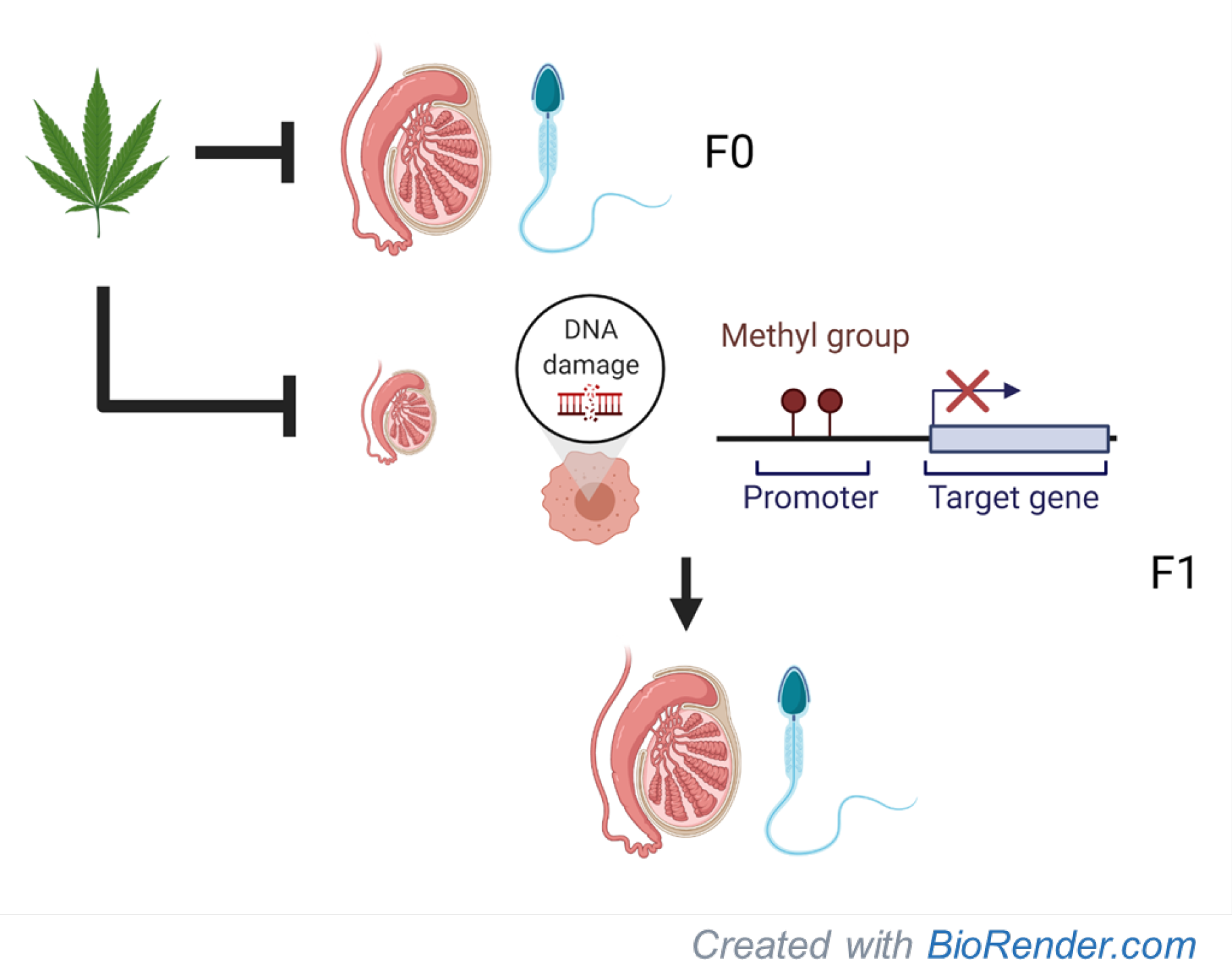

**HIGHLIGHTS:**

- Cannabis vapor exposure to adult males impairs sperm counts, motility and spermatogenesis.
- Cannabis vapor exposure to F0 males causes reduced sperm counts in F1, but not in F2 males.
- Plasma testosterone levels are not affected in F0, F1 and F2 males by cannabis exposure.
- Cannabis increases DNA damage and the expression of DNA methyltransferase (DNMT) 1 in the F1 neonatal testis.

## 1. INTRODUCTION

Cannabis is considered the most widely used recreational drug in the United States (US) and worldwide. The cannabis plants, i.e. *cannabis sativa*, can induce psychoactive effects associated with adverse mental health (Gillespie et al., 2019) and substance abuse (Patel and Marwaha, 2020). Two major components of cannabis plants, Δ^9^-tetrahydrocannabinol (THC) and cannabidiol (CBD), are well-known/well-studied phytocannabinoids, as THC is a primary psychoactive compound in cannabis plants, whereas CBD is considered to be a non or less-psychoactive compound (Maccarrone et al., 2021). In 2020, the National Survey on Drug Use and Health (NSDUH) reported that over 48 million Americans ages 12 and older used cannabis in the past year (SAMHSA, 2020). This number has been increased due to the expanding legalization of cannabis consumption in the US. Since 2012, a total of 18 states, Washington D.C. and two territories have legalized the recreational sale and use of cannabis. A total of 36 states and 4 territories allow for the medical use of cannabis products (NCSL, 2021). Following the increase in the legalized recreational cannabis use, higher potency strains of cannabis plants have emerged over the past two decades (ElSohly et al., 2016). Indeed, the average concentration of THC has increased from 9% to 17% past 10 years, and the ratio of THC and CBD also has risen substantially from 23 in 2008 and to 104 in 2017 (Chandra et al., 2019). Unsurprisingly, adverse health outcomes associated with cannabis consumption have also been increasing (Belladelli et al., 2021).

Because THC induces psychoactive effects, many studies have focused on the cannabis-induced alterations in the central nervous system (CNS). However, cannabis is also known to negatively impact reproductive functions (Hsiao and Clavijo, 2018; Payne et al., 2019; Dubovis and Muneyyirci-Delale, 2020). Importantly, many normal reproductive aspects are modulated by the endocannabinoid system (ECS). Small natural lipids, endocannabinoids such as N-arachidonoylethanolamine (anandamide; AEA) and 2-arachidonoylglycerol (2-AG), are essential regulators in reproduction (Barchi et al., 2019; Maccarrone et al., 2021), including the hypothalamus-pituitary-gonad (HPG) axis, germ cell development and sperm functions (Ricci et al., 2007; Cacciola et al., 2008; Grimaldi et al., 2013; Barchi et al., 2019), as well as their biological actions are mediated by cannabinoid receptors (CB1 or CNR1 or CB2 or CNR2). The action of THC, the exogenous property of cannabinoids, is also primarily mediated by CB1 and/or CB2 receptors (Pertwee et al., 2010). Thus, THC can negatively over-activate ECS in reproductive functions. In fact, robust reduction of sperm counts and motility associated with cannabis use have been reported in human cannabis users (Gundersen et al., 2015; Carroll et al., 2020). Bolus administration of THC causes decreased libido and sperm counts, as well as disruption of estrous cycles in rats (Dhawan and Sharma, 2003). THC interferes with the cellular functions of sperm, oocyte and embryo [reviewed in (Dubovis and Muneyyirci-Delale, 2020)]. Moreover, strong correlations exist between cannabis use by pregnant women and adverse obstetrical, pregnancy and neonatal outcomes [reviewed in (Dubovis and Muneyyirci-Delale, 2020; Fonseca and Rebelo, 2021)]. Furthermore, parental cannabis use increases the risks of congenital malformation (Williams et al., 2004; Weinsheimer et al., 2008; van Gelder et al., 2010; van Gelder et al., 2014; Reece and Hulse, 2019), psychosis (Zammit et al., 2009; Day et al., 2015; Grant et al., 2018; Fine et al., 2019), and delinquent behavior (Goldschmidt et al., 2000; Porath and Fried, 2005; Day et al., 2011; Sonon et al., 2015) among *in utero* exposed offspring. Recent studies further highlight the ability of THC exposure to induce significant changes in DNA methylomes in spermatozoa (Murphy et al., 2018). However, the long-term consequences of cannabis use on reproductive functions and how it might impact the next generation have not been examined.

In the present study, we examined the generational effects of cannabis vapor exposure on male reproductive function. Vaporization is the most common route of cannabis administration in humans (Baggio et al., 2014; Lee et al., 2016), whereas purified THC, CBD and/or synthetic CB1 agonists have been dosed to animals via either intraperitoneal injection (i.p.), intravenous exposure (i.v.) or oral gavage (p.o.) to assess the physiological and/or toxicological effects of cannabis (Carvalho et al., 2020). Approximately 120 unique phytocannabinoids are present in *Cannabis sativa* (Cather and Cather, 2020) and can vary depending on the cannabis chemovars. These facts indicate that THC or other compounds alone with direct i.p or i.v administration might not represent appropriate cannabis effects in humans. Therefore, in order to understand the generational effects of cannabis exposure on male reproductive functions, the present study was performed using an inhalation method as an administration route, by which adult male mice were exposed to dry cannabis plants to assess the toxicological effects of cannabis on F0, F1 and F2 male reproductive functions. We report that cannabis vapor exposure to adult males impairs sperm counts and/or motility and disrupts spermatogenesis in F0 and F1 males. Furthermore, cannabis increases DNA damage and the expression of DNA methyltransferase (DNMT) 1 in the F1 neonatal testis. Our results suggest that vapor exposure to cannabis generationally affects male reproductive functions, probably due to spermatogenic defects in the developing testis, potentially via altered epigenetic modification.

## 2. MATERIALS AND METHODS

### 2.1. Cannabis Vapor Exposure

The vapor chamber (Allentown Inc.) established in the Davis laboratory (Brutman et al., 2019) was constructed by attaching a commercial cannabis vaporizer (VapirRise 2.0, Vapir Inc.) to standard rodent cages (see Fig. 1A). Dry plant matter *cannabis sativa* (Code 7360, 6.7% THC, 0.02% cannabidiol, CBD) classified as “bulk marijuana” was obtained from National Institute on Drug Abuse (NIDA) Drug Supply (Research Triangle, NC). Cannabis consisted of ground whole plant matter, with large stems removed, prior to combustion in the Vaporizer. Vaporized cannabis was delivered to a positively ventilated chamber with a flow rate < 1L/min.

**Figure 1.**
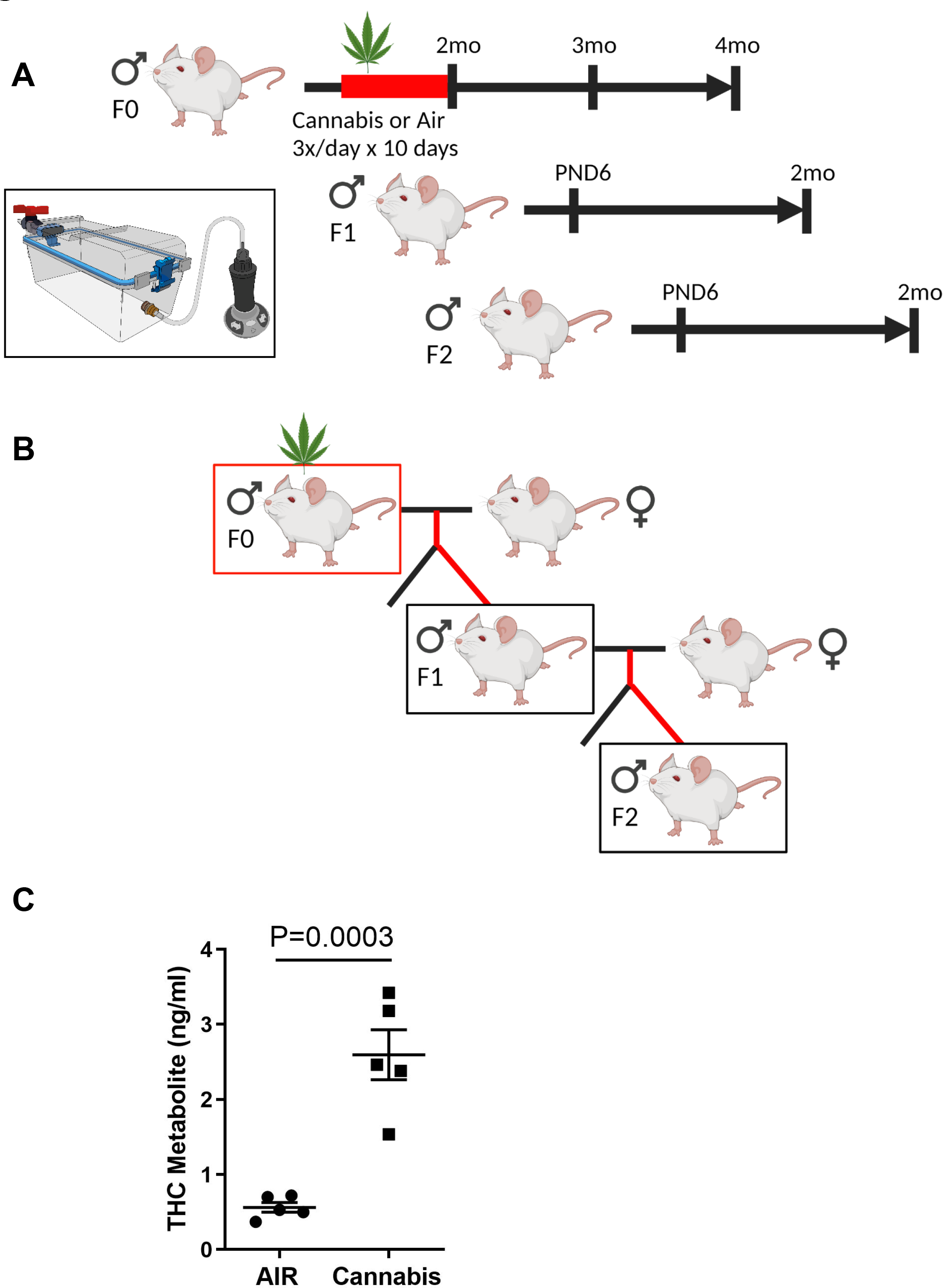
Schematic diagram of the experimental design. (A) Adult CD-1 male mice (F0) were exposed to air as a control, cannabis (200mg) 3x/day over a 10 day period and euthanized at 2-mo, 3-mo and 4-mo (n=5/group). F1 and F2 males at postnatal day (PND) 6 and 2-mo of age were also characterized. (B) F0 mice were bred to drug-naïve CD-1 females to generate F1 offspring. The breeding males (F0) were euthanized at 3-mo of age. F1 males were bred to drug-naïve CD-1 females to generate F2 offspring (n=5/group). (C) THC metabolite in plasma was analyzed 18 hr after the last cannabis exposure (n=5/group). (AB) Created with BioRender.com

### 2.2. Animals and Study Design

CD-1 male mice were purchased from Envigo (Indianapolis IN), and maintained in the vivarium at Washington State University according to institutional guidelines for the care and use of laboratory animals. A schematic outline to expose founders and generate F1 and F2 male offspring and study details are shown in Fig. 1AB. Adult male mice (6-7 weeks old, F0) were assigned based on the equal distribution of body weights to two different groups (air and cannabis, n=15 each), and mice were individually exposed to air (control) or 200 mg of vaporized cannabis plant matter for a 10 day period. To closely mimic current cannabis use in humans, F0 males were exposed to cannabis vapor at 6 am, 12 pm, and 6 pm (3x/day). Adult males exposed to air or 200 mg of cannabis 3x/day over a 10 day period displayed increased plasma levels of the primary THC metabolite, tetrahydrocannabinol carboxylic acid (THC-COOH, see details of the ELISA) at 18 hr after the last session (2.9 ± 0.3 ng/ml) relative to air-exposed controls (Fig. 1C). Note: THC-COOH is a major metabolite of THC, that has a long half-life and is generally used for the detection of cannabis use in humans. Plasma levels of THC-COOH in humans have been reported approximately 7-60 ng/ml in the frequent cannabis smoke users (Karschner et al., 2009; Schwope et al., 2011). According to allometric scaling and the FDA guideline using a standard conversion factor of 12.3 as a dose equivalence between mouse and human (Nair and Jacob, 2016), plasma levels of THC-COOH in mice is estimated 0.6-4.9 ng/ml. Thus, the detected level, approximately 3 ng/ml of THC-COOH in plasma, is considered to be the relevant comparative dose to humans. Importantly, in a dose-response of vaporized cannabis with 200 mg reliably and significantly stimulated appetite in mice. On the next day after the final vapor exposure, mice were assigned to 3 different time points (2-, 3- or 4-mo) for the sample collection. The first five males from each group (n=5/group) were euthanized to examine the direct effect of cannabis exposure on male reproductive function, as noted 2-mo. The second five males (n=5/group) were bred to drug-naïve CD-1 females to generate F1 offspring. The breeding males in the second group were euthanized at 3-mo of age. The last five males from each group (n=5/group) were kept until 4-mo of age to assess the long-term effect of cannabis exposure. One of the littermates from F1 males was bred to drug-naïve CD-1 females to generate F2 offspring.

On 2-, 3- and 4-mo (F0 males), and postnatal day (PND) 6 and 2-mo (F1 and F2 males), both testes from one male per litter (n=5 litters/group/age) were collected. One testis was cut half and homogenized with Trizol (Thermo Fisher) for RNA isolation and snap-frozen for protein extraction, and the other was fixed in Bouin’s for histological analysis. Body and paired testes weights were recorded. Blood from adult males was collected by cardiac puncture, and plasma was stored at -20°C for hormone assays. Caudal epididymis from adult males was collected for sperm count and motility assays.

### 2.3. Sperm counts

Sperm counts were performed as described previously with some modifications (Shi et al., 2017, 2018; Shi et al., 2019). Briefly, caudal epididymis was dissected and placed in 1 mL of EmbryoMax heated to 37 °C and incubated for 15 min. After the incubation, 10 µl was placed in the center of a Cell-Vu sperm counting cytometer and analyzed by SCA^®^CASA system (Fertility Techinology Resources, Inc.) under the non-capacitating conditions following the manufacturer’s instructions.

### 2.4. ELISA

Plasma levels of testosterone (T, sensitivity 90 pg/ml) and THC metabolite (sensitivity 0.072 ng/ml) were measured by ELISA (T: EIA1559, DRG Diagnostics, and THC metabolite: 701570, Cayman Chemical) following the manufacturer’s instructions. Note: THC ELISA kit indicates 100%, 28%, 18%, 15% and 12% cross-reactivities to 11-nor-9-carboxy-Δ9-THC, 11-nor-hydroxy-Δ9-THC, Δ8-THC, 11-hydroxy-Δ9-THC and Δ9-THC, respectively.

### 2.5. Quantitative real-time PCR (qPCR) analysis

Total RNA was isolated from the testis. RNA quality was assessed by spectroscopy, and the cDNA was synthesized from total RNA (1 µg) using high-capacity cDNA reverse transcription kit. QPCR was performed using StepOnePlus Real-Time PCR System (Applied Biosystems) with SYBR Green as the detector according to manufacturer’s recommendations. Primers (Supplementary Table 1) were designed to amplify exon-spanning cDNAs, and all exhibited similar amplification efficiency (95 ± 3%). Threshold cycle (CT) values were determined as the cycle number at which the threshold line crossed the linear extrapolation of the amplification curve. Background signals were established with -RT negative control templates.

After amplification, the specificity of the PCR was determined by both melt-curve analysis and gel electrophoresis. Data were normalized against *Rpl19* and are shown as the average fold increase ± standard error of the means (SEM). The fold changes are equivalent to 2^x-y^ where x is the CT value of the control and y is the CT value of treatments.

### 2.6. Immunohistochemistry and TUNEL analyses

Immunolocalization of phosphorylated forms of CB1, CB2, γH2AX, DNMT1, DNMT3A and DNMT3B was visualized in cross-section (5 µm) of paraffin-embedded tissues using specific primary antibodies and a Vectastain Elite ABC Kit (Vector Laboratories) or AlexaFluor 488-conjugated secondary antibodies (Invitrogen). Antibodies used in these analyses are listed in Supplementary Table 2. The terminal deoxynucleotidyl transferase dUTP nick end labeling (TUNEL) assay was performed according to manufacturer’s instructions using ApopTag Fluorescein In Situ Apoptosis Detection Kit (Millipore).

### 2.7. Statistical analysis

Data were analyzed by a two-tailed Student’s t-test comparing between air and cannabis group using Prism 9.1.0 (GraphPad Software). F-test was used to determine whether two groups possessed equal variances. All experimental data are presented as mean with standard error of the mean (SEM). Unless otherwise indicated, a *P* value less than 0.05 was considered to be statistically significant.

## 3. RESULTS

### 3.1. Effects of cannabis exposure on reproductive functions and spermatogenesis in F0 males

To investigate the effect of direct vapor exposure to cannabis on male reproduction, we first examined the body and testis weights, sperm counts and motility at 2-mo, a day after last cannabis exposure, and 3- and 4-mo as long-term effects of cannabis exposure in F0 males (Figs. 2 and 3). Vapor exposure to cannabis did not affect body weight, testis weight and the ratio of testis to body weight in F0 males at 2-, 3- and 4-mo (Fig. 2ABC). Mean sperm counts in the control air group did not show a significant difference in those of the cannabis group at the 2-mo of age (Fig. 3A), whereas mean sperm motility in the cannabis group (∼57%) was decreased compared with those in control (71%). In contrast, the long-term effects of cannabis at 3-mo decreased sperm counts from 18.9 ± 3.7 x 10^6^ in control to 9.1 ± 1.9 x 10^6^ in cannabis, while sperm motility was not altered in the cannabis group (Fig. 3B). Two months after the exposure at 4-mo old, sperm counts were not affected by cannabis, but sperm motility was increased in the cannabis group (Fig. 3C).

**Figure 2.**
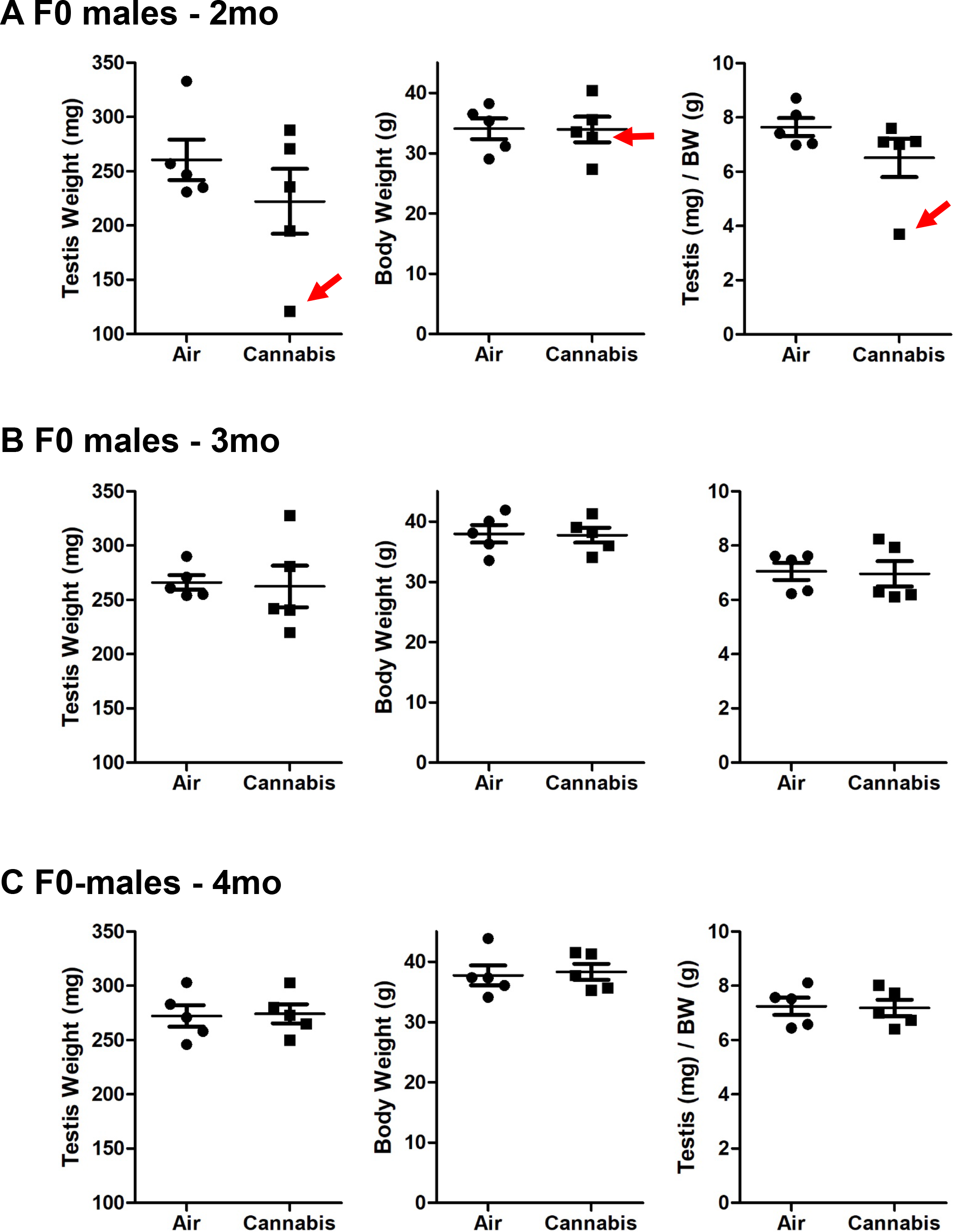
Effects of cannabis vapor exposure on testis weight, body weight, and the ratio of testis to body weight at (A) 2-mo, (B) 3-mo and (C) 4-mo F0 males (n=5/group). The red arrows show abnormal testis exposed to cannabis (see details in text and Supplementary Fig. 3).

**Figure 3.**
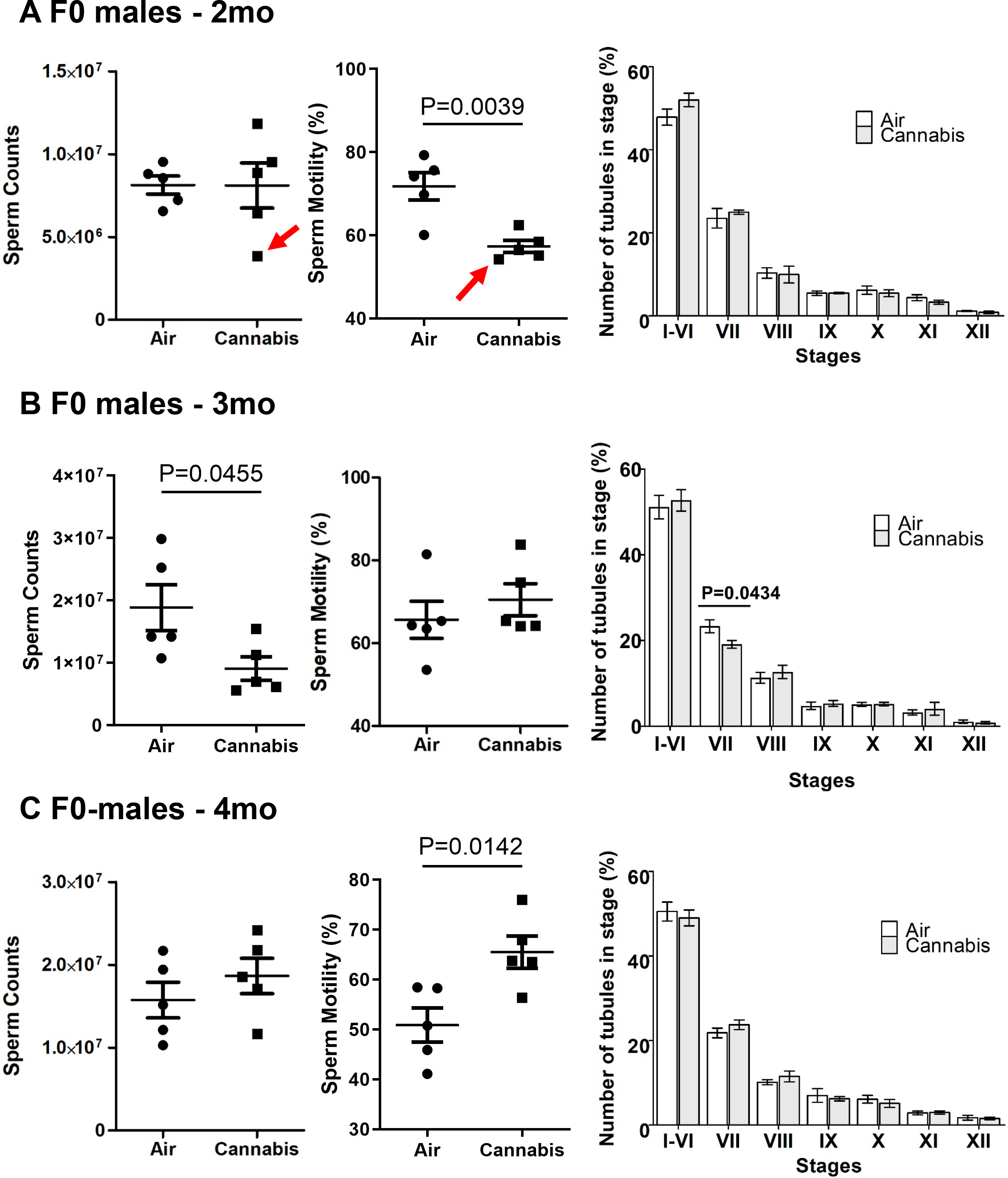
Effects of cannabis vapor exposure on sperm counts, motility and stages of the spermatogenic cycle in F0 males at (A) 2-mo, (B) 3-mo, and (C) 4-mo. Sperm counts and motility were examined from the average sperm counts of both epididymides in each animal. Comparison of the relative proportions of the 12 distinct stages of spermatogenic cycle in mice exposed to air or cannabis were examined. Only round tubules cross-section of a single, clearly defined stage were scored with the number of qualifying tubules ranging from 184 to 301 per animal (n=5/group).

The spermatogenic cycle of the seminiferous epithelium is divided into 12 stages that distinguish the progression of germ cell development. To examine whether spermatogenesis in F0 males was compromised by cannabis exposure, we quantified the distribution of testis tubule stages (Fig. 3) as described previously (Shi et al., 2017, 2018; Shi et al., 2019). Cannabis exposure decreased the proportion of tubules in stages VII in F0 males at 3-mo, although there were no significant alterations in the distribution of stages at 2- and 4-mo.

We did not observe any obvious abnormal testicular histologies in control air and cannabis males (Supplementary Fig. 1) except one cannabis exposed male, C15 (Supplementary Fig. 2). We would like to report one of the males, C15, in the cannabis group showing abnormal testis, which includes smaller testis and low sperm counts/motility (red arrows in Figs. 2A and 3A). Histological analyses indicated numerous shrunken tubules in C15 (Supplementary Fig. 2b) compared with those in control (Supplementary Fig. 2a). These were further characterized as empty seminiferous tubules with absent spermatozoa (Supplementary Fig. 2c), spermatocytes undergoing apoptosis (black arrows in Supplementary Fig. 2c), and disarranged cell layers with multi-nucliated cells (black arrows in Supplementary Fig. 2d). Generally, apoptotic cells are barely found in the adult testis, as shown in Supplementary Fig. 2e. However, TUNEL analysis revealed that there were many apoptotic cells in the testis of C15 cannabis-exposed males (Supplementary Fig. 2f). Note: Since this male showed abnormal testicular histology including undetectable stage identities due to loss of germ cells, this animal was excluded for the analysis of the stage distribution (Fig. 3A).

### 3.2. Effects of cannabis exposure on reproductive functions and spermatogenesis in F1 and F2 males

Next, to understand the generational effects of cannabis exposure on male reproductive functions, F1 and F2 offspring were examined (Fig. 4). As we expected, body and testis weights and the ratio of testis to body weights were not affected by cannabis exposure at 2-mo of F1 and F2 males (Fig. 4AB). Interestingly, we observed that sperm counts in F1 offspring were significantly decreased by cannabis exposure (Fig. 4C). However, sperm motility in F1 and F2 males and sperm counts in F2 males were not affected by cannabis (Fig. 4CD). Furthermore, there were no significant differences in the distribution of seminiferous epithelium stages (Fig. 4CD).

**Figure 4.**
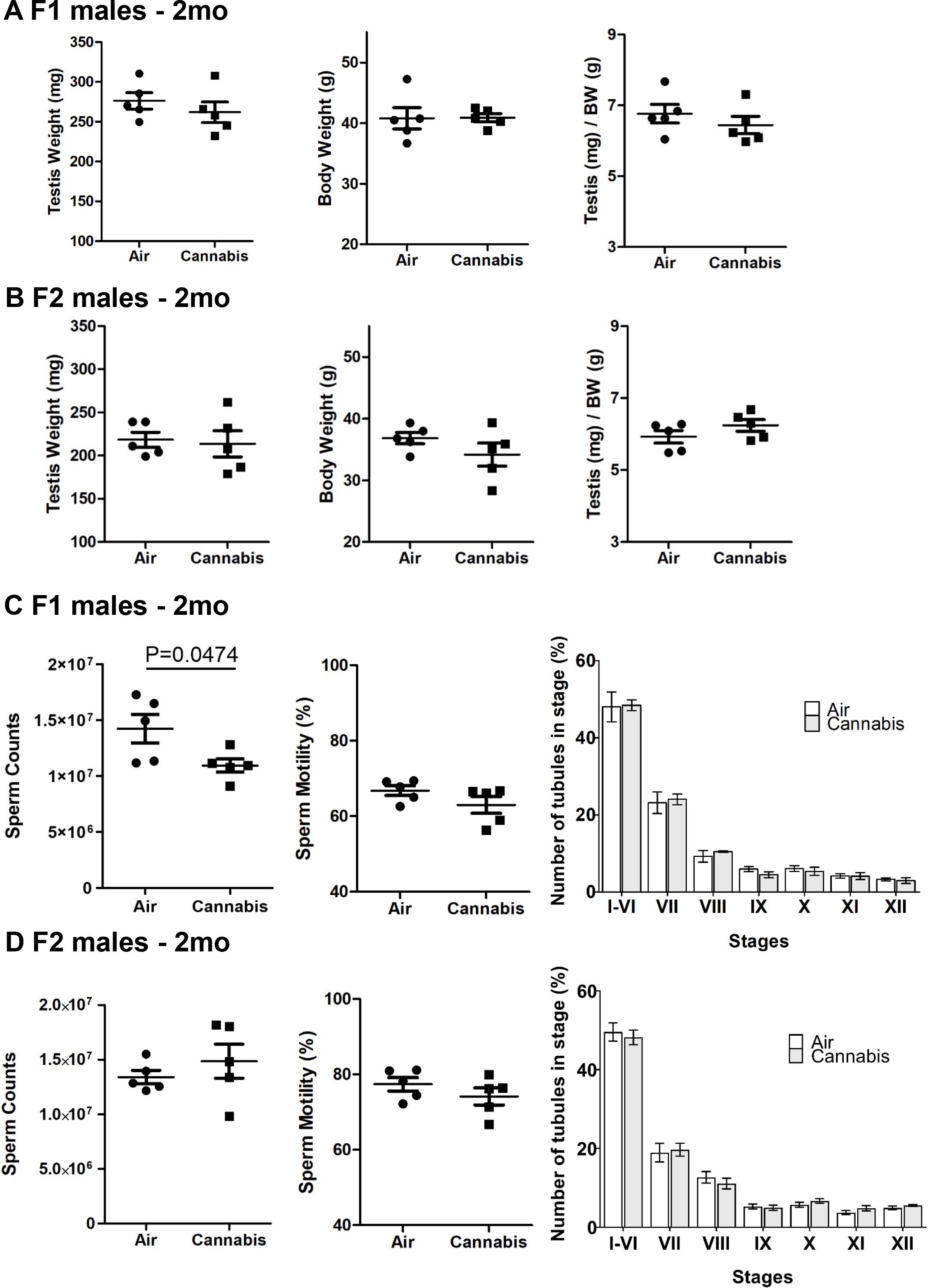
Effects of cannabis vapor exposure on F1 and F2 males (n=5/group). (A) Testis weight, body weight and the ratio of testis to body weight in F1 males. (B) Testis weight, body weight and the ratio of testis to body weight in F2 males. (C) Sperm counts, motility and stages of the spermatogenic cycle in F1 males. (D) Sperm counts, motility and stages of the spermatogenic cycle in F2 males.

### 3.3. Effects of cannabis exposure on cannabinoid receptor expression and steroidogenesis

ECS via cannabinoid receptors such as CB1 or CNR1 and CB2 or CNR2 regulates Leydig cell development, germ cell differentiation and sperm functions in male reproduction (Ricci et al., 2007; Cacciola et al., 2008; Grimaldi et al., 2013; Cobellis et al., 2016). It is also known that THC activities are mainly mediated by cannabinoid receptors (CB1 and CB2) (Pertwee et al., 2010). We thus sought to examine the expression of CB1 and CB2 receptors in the testis of air and cannabis exposed animals. Immunohistochemical analysis showed CB1 was specifically localized in the Leydig cells in the adult testis (Fig. 5A), whereas we could not detect CB2 using a commercially available antibody. *Cnr1* mRNA was detectable in the adult testis; however, *Cnr2* was below the detection level by QPCR using two different sets of primers, supporting undetectable immunoreactive CB2 in the adult testis. *Cnr1* mRNA levels were not altered in the testis from F0 mice at 2-mo, 3-mo and 4-mo and in F1/F2 mice at 2mo, while its level in the F0 3-mo testis tended to be increased by cannabis exposure (Fig. 5B), but did reach significance (P=0.136).

**Figure 5.**
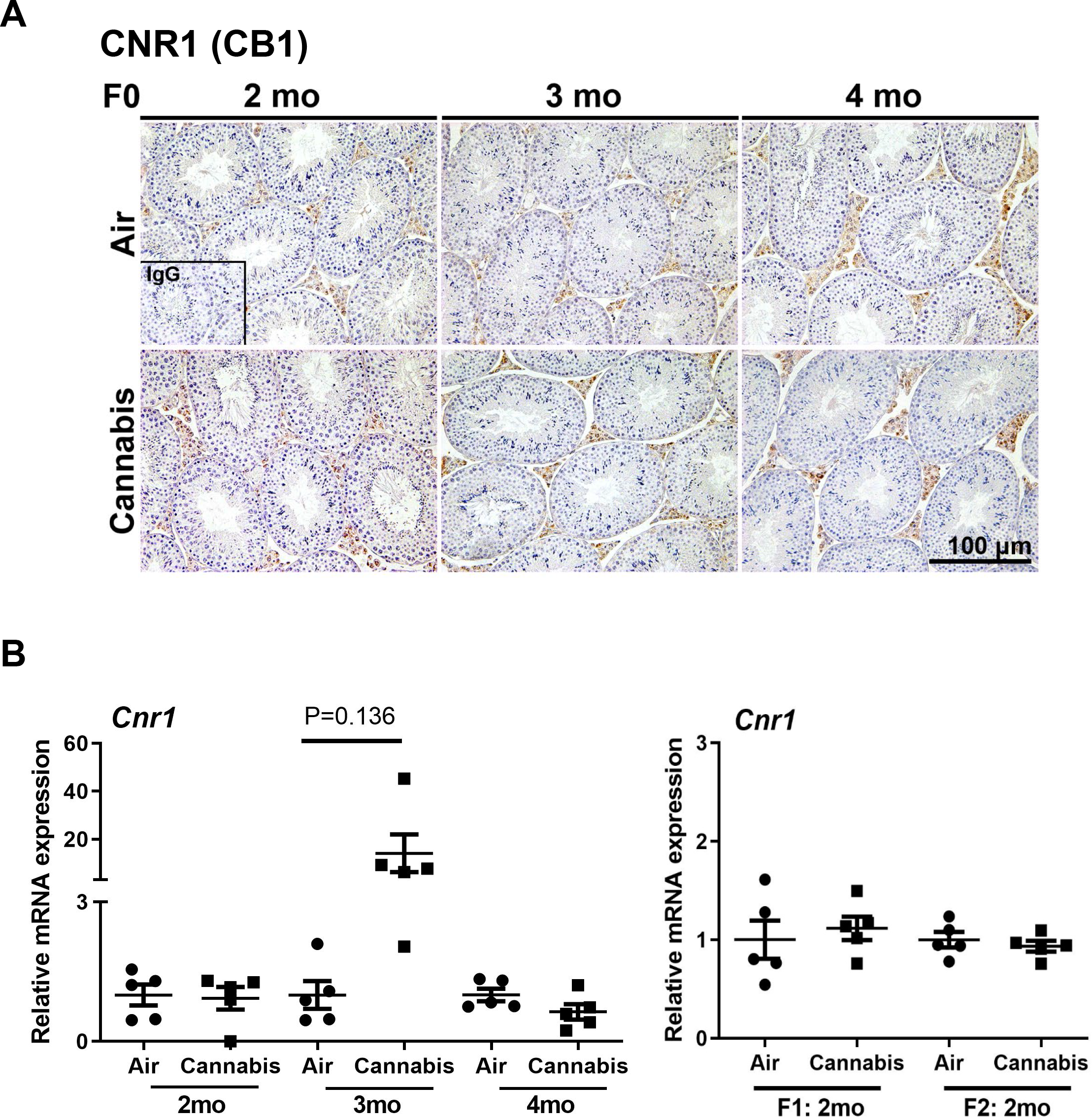
Effects of cannabis vapor exposure on CB1 receptor expression analyzed by (A) immunohistochemistry and (B) QPCR in adult F0, F1 and/or F2 testis (n=5/group).

As CB1 was strongly expressed in Leydig cells, where testosterone is mainly produced, plasma testosterone levels and steroidogenic enzymes in the testis were assessed to determine whether cannabis exposure affected reproductive hormone levels and their synthesis. Although testosterone levels in the plasma were not altered by cannabis exposure (Fig. 6A), the relative mRNA expressions of *Cyp11a1* at 2-mo and *Cyp19a1* at 3-mo in F0 males were increased in the testis (Fig. 6B).

**Figure 6.**
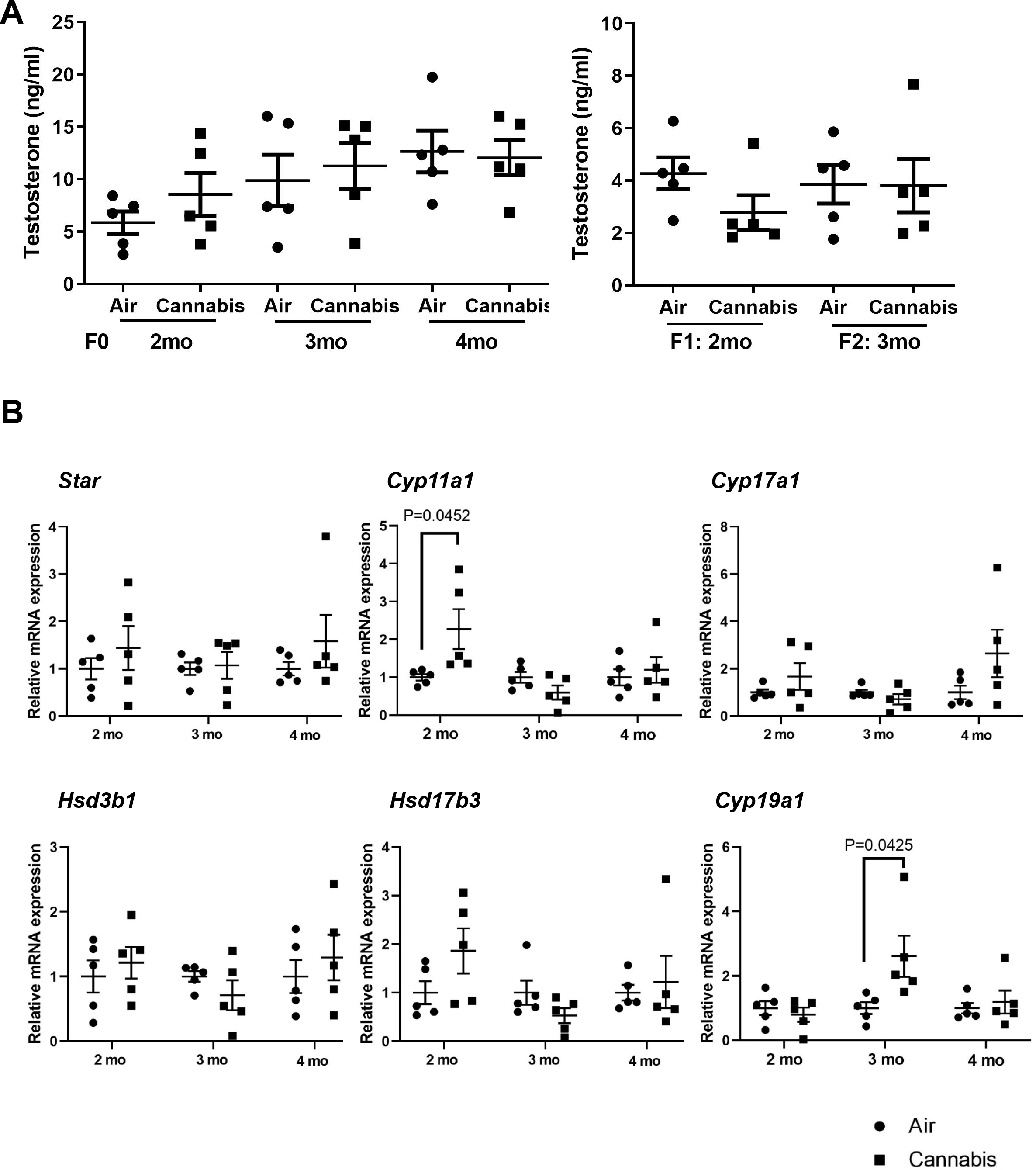
Effects of cannabis vapor exposure on steroidogenesis. (A) Testosterone levels in plasma from F0, F1 and F2 males were analyzed by ELISA (n=5/group). (B) Relative mRNA expression levels of steroidogenic enzymes in the F0 testis were analyzed by qPCR (n=5/group). The results were normalized against *Rpl19* and expressed as fold above control treatment, which was arbitrarily set to 1.

### 3.4. Effects of cannabis exposure on DNA damage and apoptosis in the neonatal testis

As cannabis exposure reduced sperm counts in F1 males (Fig. 4), we sought to determine whether spermatogenesis and testis development were altered by examining DNA damage and apoptosis. Immunostaining for phosphorylation of γH2AX revealed that cannabis exposure increased % of positive tubules and positive cells per tubule in F1 neonatal testis on PND6 (Fig. 7A), whereas TUNEL-positive cells were invariant between control air and cannabis groups (Fig. 7B). Note: We did not observe any differences between F1 and F2 offspring for litter size, sex ratio, testis weight, body weight and the ratio of testis to body weight in the neonates, except that cannabis exposed F1 neonates showed a reduced ratio of testis to body weight (Supplementary Fig. 3)

**Figure 7.**
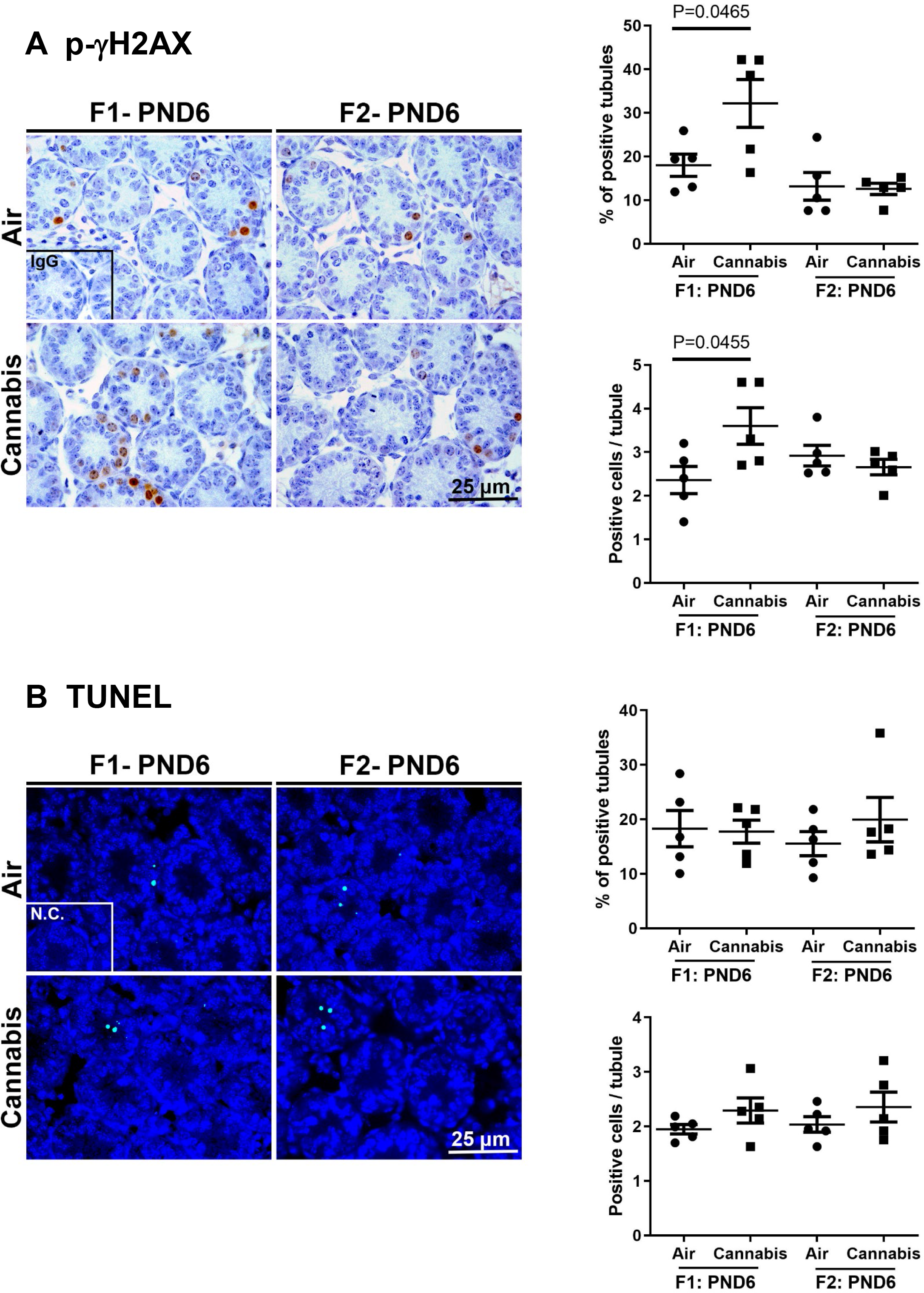
Effects of cannabis vapor exposure on (A) DNA damage and (B) apoptosis in the neonatal testis on postnatal day (PND) 6 from F1 and F2 males (n=5/group). DNA damage was assessed by immunostaining for phosphorylation of γH2AX. Cellular apoptosis was determined by TUNEL analysis.

### 3.5. Effects of cannabis exposure on DNA methyltransferases (DNMTs) in the neonatal testis

Owing to epigenetic programming is necessary for spermatogenesis in the developing testis (Godmann et al., 2009; Trasler, 2009), three active mammalian DNA methyltransferases, DNMT1, DNMT3A and DNMT3B were examined in F1 and F2 neonatal testis (Fig. 8AB). We have previously reported that DNMT1 is specifically positive in neonatal germ cells co-staining with DDX4 as a germ cell marker and GATA4 as a Sertoli cell marker (Shi et al., 2019). DNMT3A is expressed in peritubular myoid cells (PTM) and Sertoli cells, and DNMT3B is detected in germ cells (Shi et al., 2019). The present results confirmed that a few DNMT1 positive germ cells were observed in the air control neonatal testis. In contrast, cannabis-exposed F1 mice showed increased numbers of DNMT1 positive germ cells in the testis (Fig. 8A). However, we did not observe the differences in DNMT1 expression in F2 neonatal testis (Fig. 8A). Fig. 8B shows positive DNMT3A in Sertoli cells and PTM. However, its expression level was not affected by cannabis exposure in F1 and F2 neonatal testis (Fig. 8B). DNMT3B was detected in germ cells, but no differences were observed in either groups or generations (Fig. 8B).

**Figure 8.**
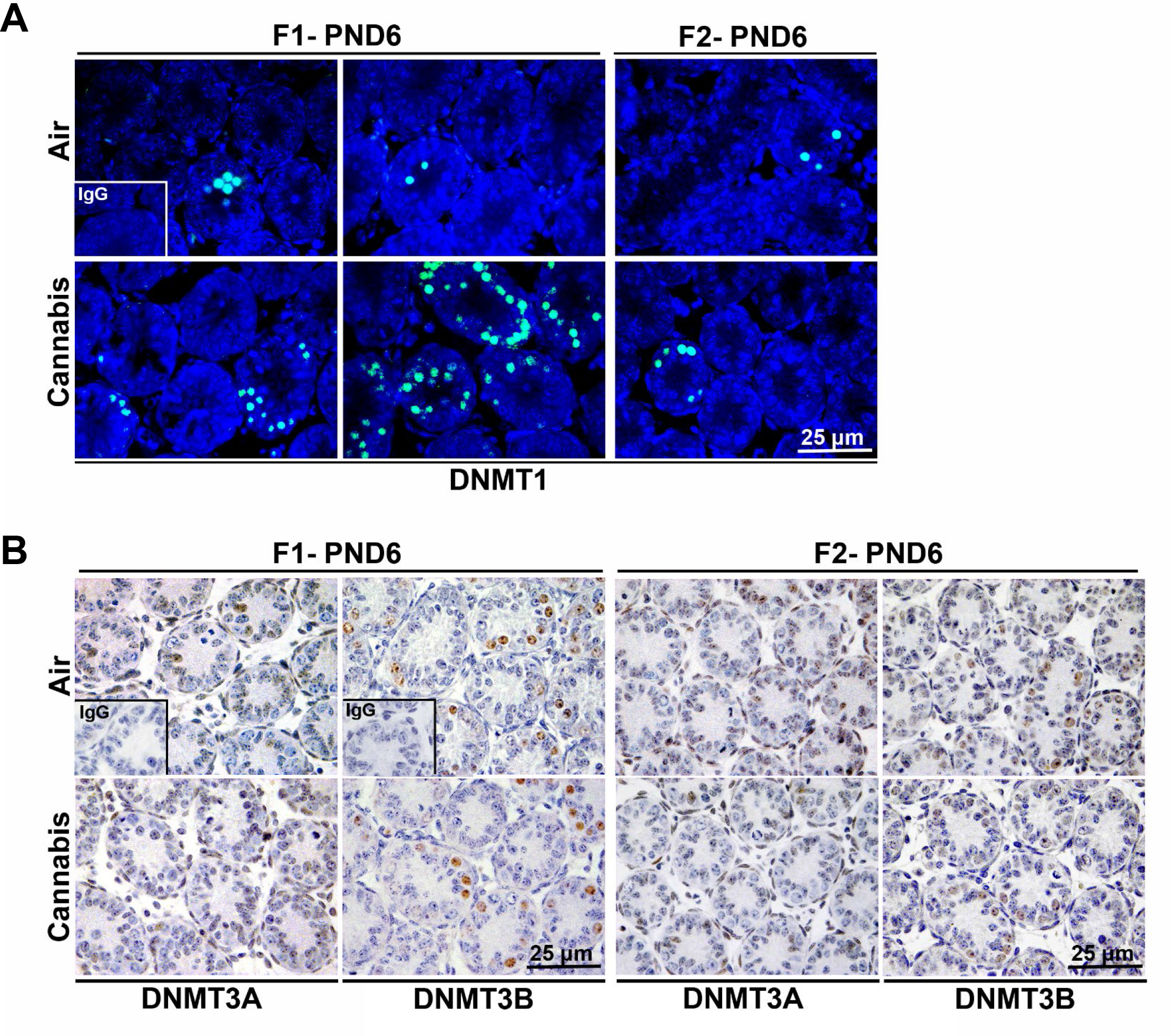
Effects of cannabis vapor exposure on DNMTs in the neonatal testis. Representative image of immunolocalization for (A) DNMT1, (B) DNMT3A and DNMT3B on postnatal day (PND) 6 (n=5/group).

## 4. DISCUSSION

The present study showed that cannabis vapor exposure in adult male mice induced a subsequent reduction of sperm counts and/or motility in F0 and F1 males. Altered and disrupted spermatogenic progression in F0 and F1 testis were also observed. Until recently, studies describing the effects of cannabis have relied on administering THC to rodents (Williams et al., 1998; Williams and Kirkham, 2002; Manwell et al., 2014a; Manwell et al., 2014b). However, dry cannabis plant matter, which contains numerous cannabinoid compounds (Lafaye et al., 2017), is the most commonly used form of the drug (Grant et al., 2012; Baggio et al., 2014; Lafaye et al., 2017), and vaporization is the preferred route of administration (Earleywine and Barnwell, 2007; Baggio et al., 2014; Lee et al., 2016). Thus, this is an ideal administrative method to study cannabis exposure on reproductive functions, and we believe this is the first time reporting a systematic study examining reproductive phenotypic defects and their associated mechanisms in mice using vaporization as a cannabis administration.

In the present study, sexually mature adult males were exposed to cannabis vapor. Male mice reach sexual maturity around 6 weeks of age, in which sperm appear in the epididymis after the first wave of spermatogenesis completion (McCarrey, 2013). Our results showed that cannabis exposure in adult males immediately affected sperm motility, supporting the negative impact of cannabis on sperm functions (Dubovis and Muneyyirci-Delale, 2020; Maccarrone et al., 2021). While cannabis disrupted sperm motility immediately after the exposure, sperm counts were not affected. Instead, sperm counts were decreased one month after the cannabis exposure. This result suggests that cannabis negatively impacts the spermatogenesis cycle in adult males, as the differentiation process of spermatogenesis from spermatogonia to mature spermatozoa takes ∼35 days in mice (Oakberg, 1956). In fact, when we examined the stages of spermatogenesis, cannabis exposure altered stage VII. However, these effects might not persist with the next wave of spermatogenesis, as 4-mo old males, 2-mo after cannabis exposure, did not show any reduction of sperm counts and stages, although sperm motility increased.

The studies from the 1970’s have reported that direct THC i.p. administration decreases testosterone biosynthesis in rats (List et al., 1977), and THC exposure in rat Leydig cells inhibits steroidogenesis in vitro (Jakubovic et al., 1979). ECS via CB1 receptor regulates Leydig cell development and functions (Cacciola et al., 2008; Cobellis et al., 2016). However, plasma levels of testosterone were not altered by cannabis exposure in this study, although increased *Cyp11a1* and *Cyp19a1* expression in the testis was observed, suggesting that disruption of spermatogenesis and male functions might not be mainly due to altered steroidogenesis and/or inhibition of testosterone.

We used cannabis exposed males collected at 3-mo of age for the generation of F1 offspring. Examining the effects of cannabis on not only exposed males (F0) but also their offspring (F1 and F2) is crucial to understand whether phenotypic defects associated with cannabis use can be paternally transmitted to offspring. The mechanisms of how cannabis use affects the health outcomes for the next generation can be explained by epigenetic modification. Recent studies have demonstrated that parental THC exposure (Szutorisz et al., 2014; Watson et al., 2015) and paternal exposure of synthetic cannabinoid, WIN55,212-2, a CB1 receptor agonist (Ibn Lahmar Andaloussi et al., 2019) create phenotypic behavior defects correlated with altered DNA methylomes in the brain of F1 rat offspring. Studies from Murphy and colleagues highlight the ability of THC exposure to induce significant changes in DNA methylomes in the rat sperm (Murphy et al., 2018; Schrott and Murphy, 2020; Schrott et al., 2020). Furthermore, many “THC target genes” are differentially methylated in both rat sperm and brain (Watson et al., 2015; Murphy et al., 2018; Schrott et al., 2020), suggesting that paternal reproductive risks could be associated with abnormal behavior via epigenetic modification in sperm. This group has also demonstrated that DNA methylomes in human sperm are altered in cannabis users compared with those in non-user controls (Murphy et al., 2018). The present study showed increased DNMT1 expression in neonatal testis. Thus, phenotypic defects such as reduced sperm counts and/or motility in F1 males might be consequently induced by altered DNA methylation during fetal and/or neonatal spermatogenesis. In support of this, alterations of epigenetic programming of the germline during fetal development and early postnatal life disrupt male reproductive functions and promote infertility (Guerrero-Bosagna and Skinner, 2014; Nilsson and Skinner, 2015; Shi et al., 2019).

It has been considered that when the F0 father is exposed to an adverse stimulus, his child (F1) can be affected as a consequence of direct exposure of his germline to the same stimulus. The effects seen in the F2 generation that had no direct exposure to the original stimulus would be transgenerational (Xin et al., 2015). However, the present study revealed that cannabis exposure did not alter sperm counts and motility and expression of DNMTs in F2 males. These results suggest that temporal cannabis exposure to mature adult males for only 10 days might not be enough to disrupt spermatogenesis transgenerationally transmitting to the germlines since critical time points of germ cell development and spermatogenesis have been completed by 6-week-old. Thus, reduction of sperm counts and increased DNA damage and DNMT1 expression in F1 offspring are indicated due to direct exposure of cannabis to F0 males. On the other hand, accelerating cannabis use in adolescents and young adults, including pregnant women, has been reported last decades all over the world (Tirado-Munoz et al., 2020). To understand the further toxic impact of cannabis on male reproductive functions, studies of generational and transgenerational effects of prenatal and/or adolescent cannabis vapor exposure on epigenetic modifications are required and are currently underway.

## 5. CONCLUSIONS

The present study showed that cannabis vapor exposure to adult males for 10 days impaired sperm counts and/or motility and disrupted the spermatogenesis in F0 and F1, but not F2 males. Cannabis exposure increased DNA damage and the expression of DNMT1 in the F1, but not in the F2 neonatal testis. Our results indicate that vapor exposure to cannabis generationally affects male reproductive functions that can be due to spermatogenic defects in the developing testis potentially via altered epigenetic modification.

## Credit author statement

Conceptualization: Kanako Hayashi, Mingxin Shi, Emma C. Wheeler and Jon F. Davis. Experimental section: Mingxin Shi, Esther M. Langholt, Logan C. Butler, Liang Zhao, Madeleine E. Harvey, Emma C. Wheeler, Yeongseok Oh, James A. MacLean II, Emily Sabrowsky, Sue Yu, Shane Watson and Kanako Hayashi. Data analysis: Mingxin Shi, Esther M. Langholt, Logan C. Butler and Liang Zhao. Writing and Supervision: Kanako Hayashi. Writing-review and editing: Kanako Hayashi, James A. MacLean II, Jon F. Davis.

---

**Supplementary Figure 1.**
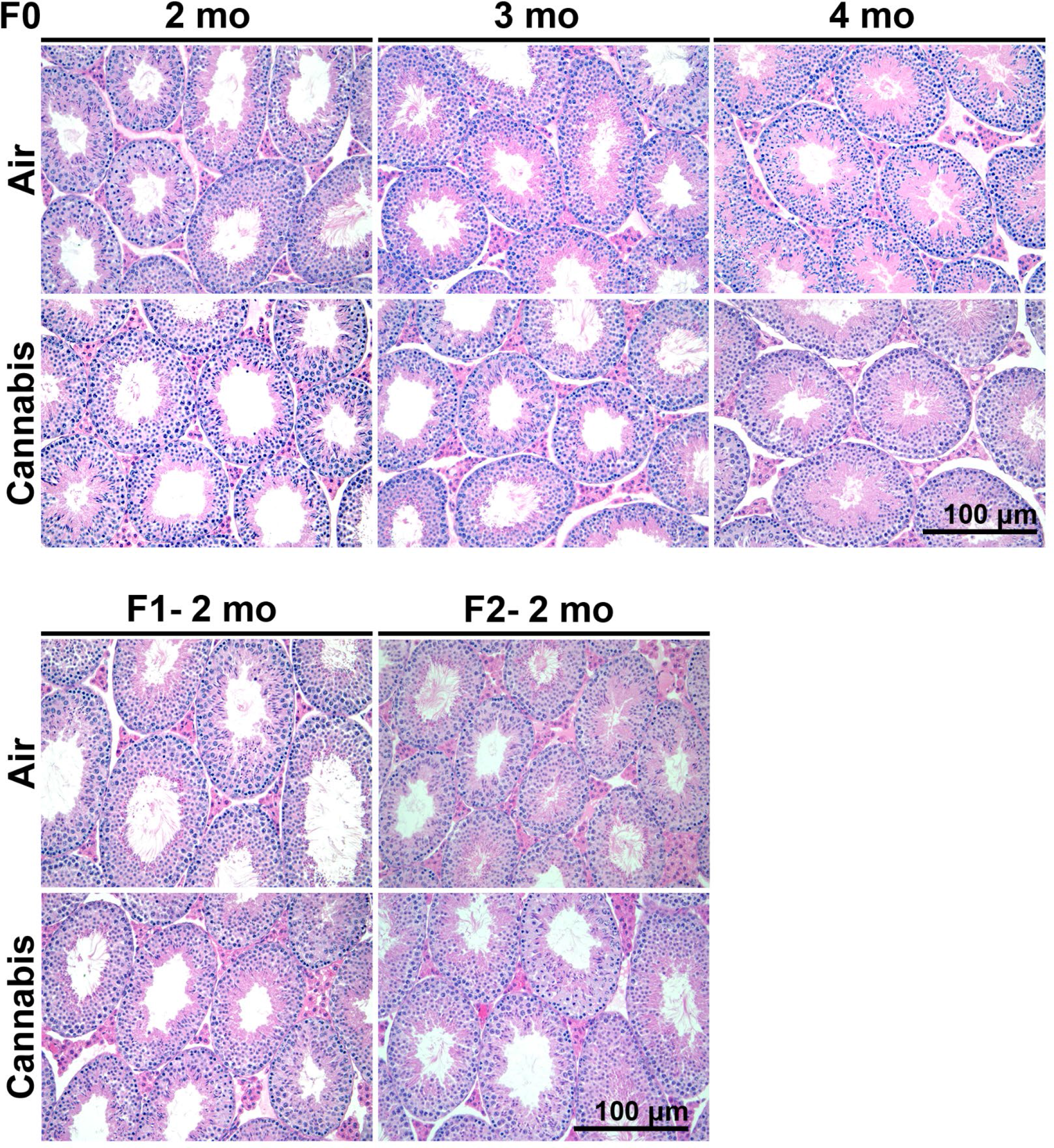
Representative histology of adult testis from F0 (2-mo, 3-mo and 4-mo), and F1 and F2 at 2mo of age.

**Supplementary Figure 2.**
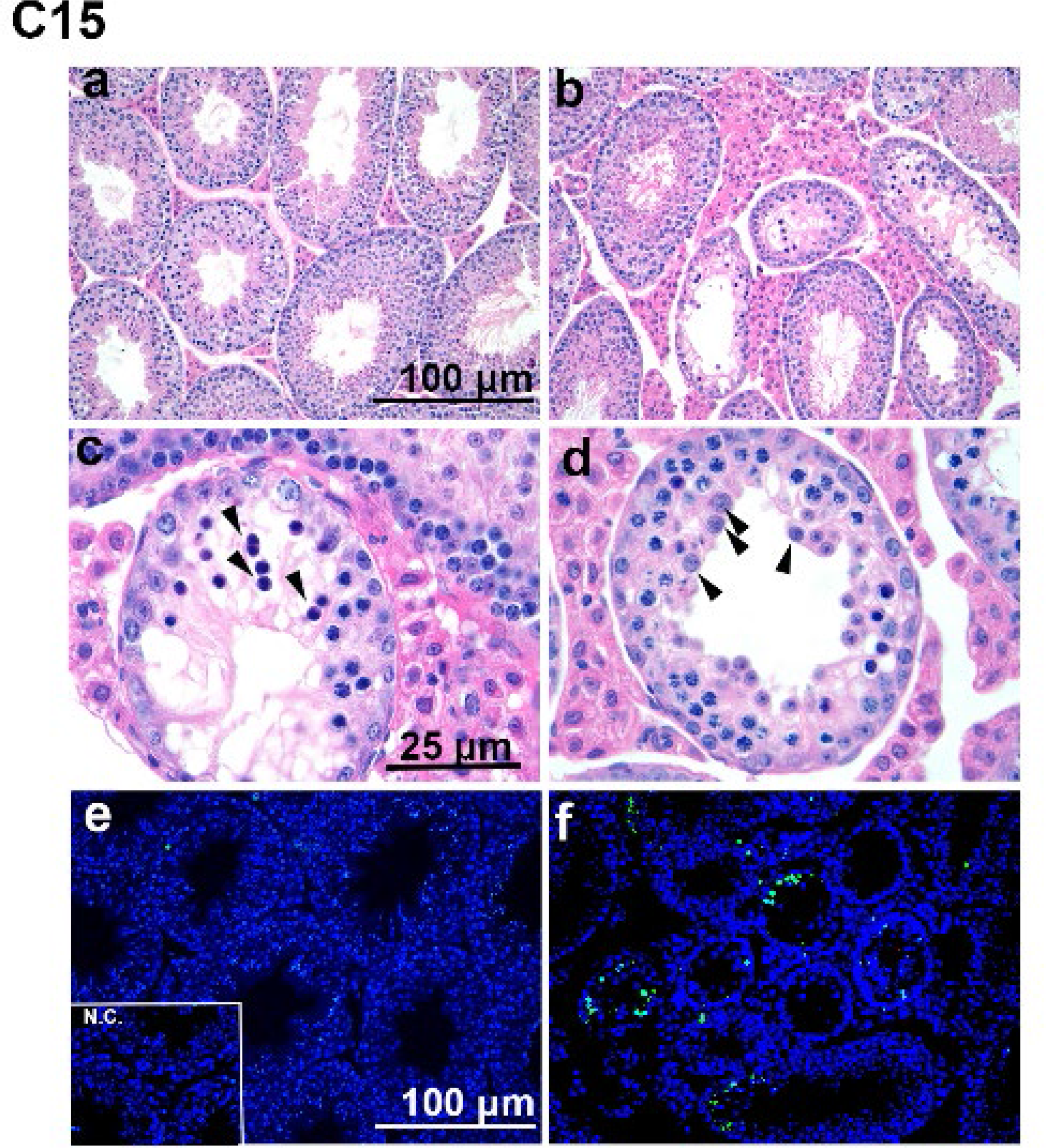
Histology of abnormal testis from control and C15, one of the cannabis exposed males. (a) Normal testis histology in control male, (b) shrunken tubules, (c) empty seminiferous tubules with absent spermatozoa, and spermatocytes undergoing apoptosis (black arrows), (d) disarranged cell layers with multi-nucliated cells (black arrows), (e) TUNEL staining in control testis, (f) TUNEL staining in C15 testis showing in many apoptotic cells.

**Supplementary Figure 3.**
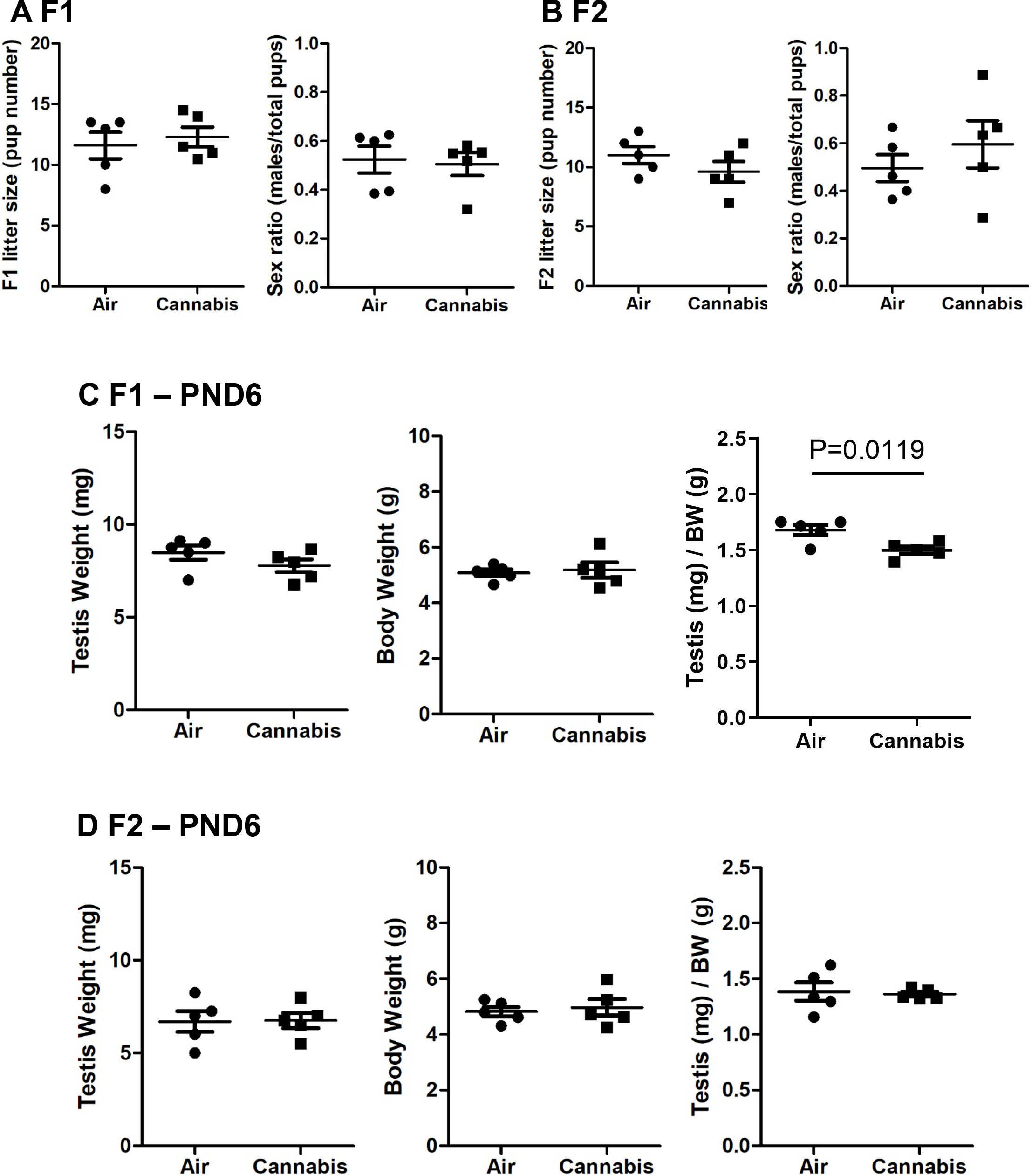
Effects of cannabis vapor exposure on (A) litter size and sex ratio of pups in F1, (B) litter size and sex ratio of pups in F2, (C) testis weight, body weight and the ratio of testis to body weight in F1 neonatal males, and (D) testis weight, body weight and the ratio of testis to body weight in F2 neonatal males (n=5/group).

**Supplementary Table 1.**
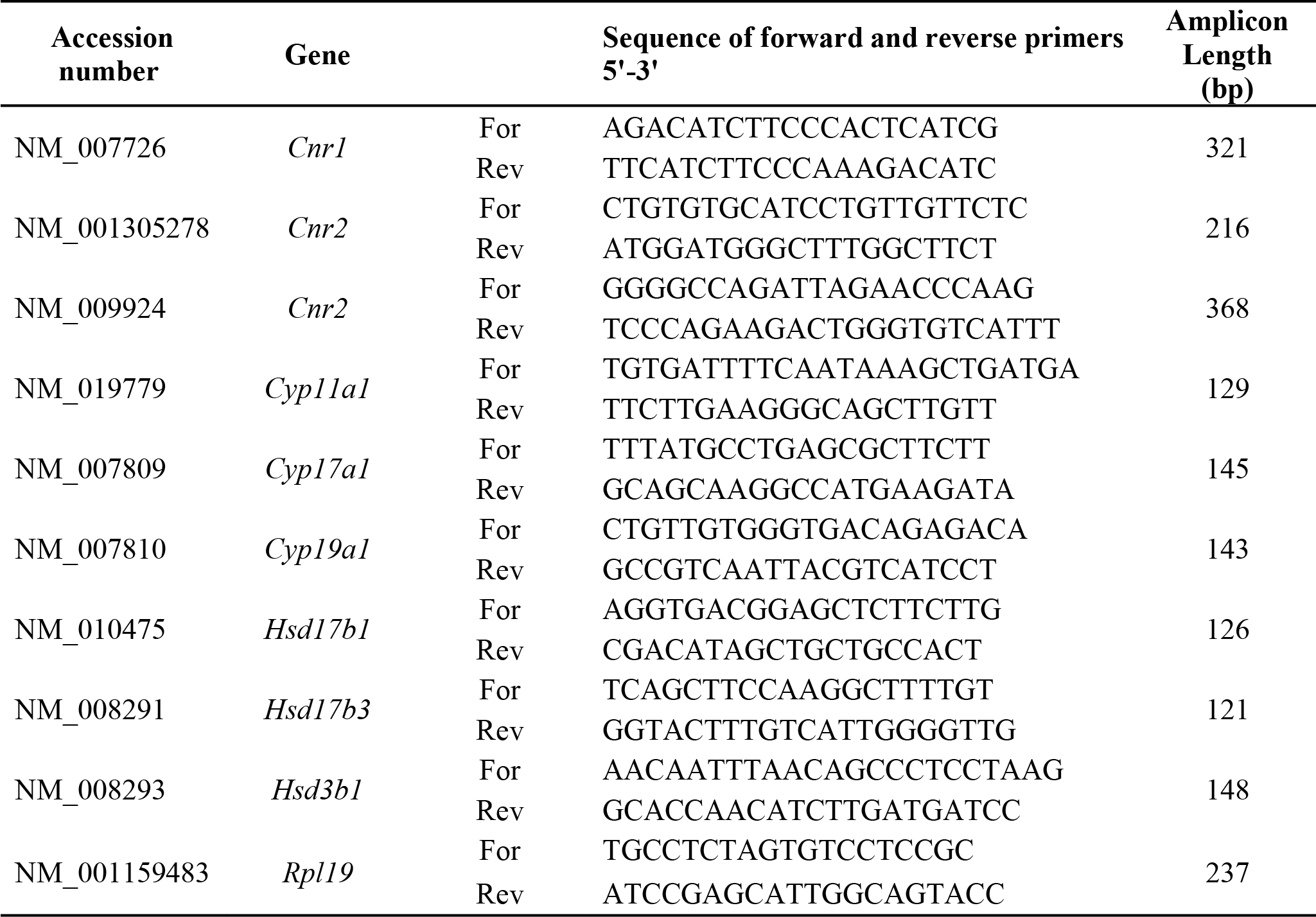
Primers for QPCR.

**Supplementary Table 2.** List of primary antibodies used for immunohistochemistry.

| **Antibody** | **Source** | **Host** | **Dilution** |
| --- | --- | --- | --- |
| CB1 | Invitrogen (PA1-743) | Rabbit monoclonal | 1:300 |
| CB2 | Abcam (ab19905) | Rabbit polyclonal | 1:50 |
| DNMT1 | Abcam (ab3561) | Rabbit polyclonal | 1:200 |
| DNMT3A | Abcam (ab2850) | Rabbit polyclonal | 1:100 |
| DNMT3B | Abcam (ab2851) | Rabbit polyclonal | 1:100 |
| Phospho-H2AX | Millipore (05-636) | Rabbit polyclonal | 1:300 |

